# GWAS associated Variants, Non-genetic Factors, and Transient Transcriptome in Multiple Sclerosis Etiopathogenesis: a Colocalization Analysis

**DOI:** 10.1101/2021.03.12.434773

**Authors:** Renato Umeton, Gianmarco Bellucci, Rachele Bigi, Silvia Romano, Maria Chiara Buscarinu, Roberta Reniè, Virginia Rinaldi, Raffaella Pizzolato Umeton, Emanuele Morena, Carmela Romano, Rosella Mechelli, Marco Salvetti, Giovanni Ristori

## Abstract

A clinically actionable understanding of multiple sclerosis (MS) etiology goes through GWAS interpretation, prompting research on new gene regulatory models. We previously suggested a stochastic etiologic model where small-scale random perturbations could reach a threshold for MS development. The recently described mapping of the transient transcriptome (TT), including intergenic and intronic RNAs, seems appropriate to verify this model through a rigorous colocalization analysis. We show that genomic regions coding for the TT were significantly enriched for MS-associated GWAS variants and DNA binding sites for molecular transducers mediating putative, non-genetic, etiopathogenetic factors for MS (e.g., vitamin D deficiency, Epstein Barr virus latent infection, B cell dysfunction). These results suggest a model whereby TT-coding regions are hotspots of convergence between genetic ad non-genetic factors of risk/protection for MS, and plausibly for other complex disorders. Our colocalization analysis also provides a freely available data resource (www.mscoloc.com) for future research on MS transcriptional regulation.

## Introduction

A large body of literature agrees that regulatory genomic intervals, especially those encompassing enhancers, are enriched with disease-associated DNA elements. Most of this evidence comes from genome wide association studies (GWAS) based on single polymorphism nucleotides (SNPs) representing common variants (Ernst et al., 2011; Farh et al., 2015; Gusev et al., 2014; Maurano et al., 2012; Vahedi et al., 2015), even though a recent study showed that low-frequency and rare coding variants may somewhat contribute to multifactorial diseases ( & Consortium, 2020). Several characteristics of regulatory disease-associated genetic variants complicate GWAS interpretation, prompting research on new gene regulatory models: (*i*) SNPs are chosen as haplotypes to spare the genotyping work needed for the large number of samples used in GWAS, therefore fine mapping and epigenetic studies are required to integrate GWAS data (Calderon et al., 2019; Mumbach et al., 2017; Ohkura et al., 2020; van Arensbergen et al., 2019); (*ii*) a fraction of supposedly causal disease-associated variants directly alters recognizable transcription factor binding motifs as it might be expected, according to their regulatory function (Farh et al., 2015); (*iii*) the identified GWAS signals are likely to exert highly contextual (i.e., time- and position-dependent) regulatory effects, that may change according to the tissue and to the time when they receive an input from inside or outside the cell. In summary, current gene regulatory models help only in part to fully detail which disease-associated SNP signals are causal, and by which exact mechanisms they are causal. Recent studies on the biological spectrum of human DNase I hypersensitive sites (DHSs), that are disease-associated markers of regulatory DNA, may help to better rework GWAS data and particularly to contextualize the genomic variants according to tissue/cell states and to gene body colocalization of DHSs (Meuleman et al., 2020). In this context, the latest version of the Genotype-Tissue Expression project may provide further insights into the tissue specificity of genetic effects, supporting the link between regulatory mechanisms and traits or complex diseases (Consortium, 2020).

Another layer of complexity comes from our recent studies suggesting an MS etiologic model where stochastic phenomena (i.e., random events not necessarily resulting in disease in all individuals) may contribute to the disease onset and progression. This model, embedded between physics of stochastic systems and cell biology, suggests how small-scale random perturbations would impact on large-scale phenomena, such as exceeding the threshold for MS development that is set by genetic and non-genetic susceptibility factors (Bordi et al., 2014; Bordi et al., 2013). Such model is consistent with our previous results on the heterogeneity of MS etiology components in twin pairs studies (Fagnani et al., 2015; Ristori et al., 2006) and with prior bioinformatics analyses that determined a significant enrichment of binding motifs for Epstein-Barr virus (EBV) nuclear antigen 2 (EBNA2) and vitamin D receptor (VDR) in genomic regions containing MS-associated GWAS variants (Ricigliano et al., 2015). We also demonstrated that genomic variants of *EBNA2* resulted to be MS-associated (Mechelli et al., 2015), and other groups expanded our findings showing that enrichment of EBNA2-binding regions on GWAS DNA intervals is involved in the pathogenesis of autoimmune disorders, including MS (Harley et al., 2018).

A recent sequencing innovation (namely, TT-seq) allowed to map the transient transcriptome that has a typical half-life within minutes, compared to stable RNA elements, such as protein-coding mRNAs, long-noncoding RNAs, and micro-RNAs, that persists at least a few hours (Michel et al., 2017; Schwalb et al., 2016; Villamil et al., 2019). The transient transcriptome (TT) includes mostly enhancer RNAs (eRNA), short intergenic non-coding RNAs (sincRNA) and antisense RNAs (asRNA). Overall, these transient RNAs (trRNA) are relatively short in length, generally lack a secondary structure, and would not present those chemical modifications that characterize unidirectional and polyadenylated stable RNAs (Natoli & Andrau, 2012; Schwalb et al., 2016).

Other recent works based on time-resolved analysis, agree on the eRNAs very rapid functional dynamics model while interacting with the transcriptional co-activator acetyltransferase CBP/p300 complex (Bose et al., 2017; Weinert et al., 2018). This confirms the highly contextual role of eRNAs through the control of transcription burst frequencies, which are known to influence cell-type-specific gene expression profiles (Larsson et al., 2019). Along these lines, a recent study showed that T cells selectively filter oscillatory signals within the minute timescale (O’Donoghue et al., 2021), further supporting the aforementioned model.

In summary, on the basis of our previous research (i.e., the heterogeneity in the MS etiology components; the stochasticity in the interaction between genetic and non-genetic factors contributing to disease development; the enrichment of binding sites for environmental factors in MS-associated DNA intervals) and leveraging the recent sequencing innovations in the mapping of the transient transcriptome (i.e., the erratic time dynamics and the highly contextual expression) (Michel et al., 2017; Schwalb et al., 2016), we hypothesize that MS-associated GWAS signals prevalently fall within regulatory regions of DNA coding for trRNAs. In theory, the genomic intervals coding for this transient transcriptome may be the hotspots where temporospatial occurrences (stochastic in nature, as said) may coalesce and so contribute to physiological (developmental and/or adaptive) outcomes, or possibly give rise to disease onset or progression. This study is aimed at verifying this working hypothesis through a colocalization analysis and its further dissection in the context of MS.

## Results

### MS-associated GWAS signals colocalize with regulatory regions of DNA plausibly coding for trRNAs

We set up our region-of-interest (ROI) inside GWAS catalogue (Buniello et al., 2019) by considering all MS GWAS that were published, extracting all SNP positions, and creating a single set of genomic coordinates that therefore encompass all GWAS-derived or GWAS-verified signal for MS. We then refined the SNP list by pruning out about 1.5% of the SNPs as they did not contain intelligible genomic annotations or were duplicates. The final ROI list is reported in Supplementary Table S1 and consists of 603 unique single-nucleotide regions; to provide a “threshold” against which the match ROI<>Database would be benchmarked, we used 107,423 regions as Universe, that corresponded to the signals coming from the entire GWAS Catalog.

Next, we matched through colocalization analyses our ROI with lists of regions resulting from the work by Michel et al., which mapped the transient and stable transcriptome captured by TT-seq after T cell stimulation (Schwalb et al., 2016). We found a significant enrichment of MS-associated genetic variants in the transient transcriptome (p-value=2.80 × 10^−9^; Table 1). Of note, when we split the transcriptome list in two subsets for long (≥ 60 minutes) and short (< 60 minutes) half-life, we found that only the short half-life subset significantly colocalized with the ROI (p-value 2.06 × 10^−8^ vs 0.09). This finding was indicative of the relationship between MS-associated GWAS signals and the regulatory regions of DNA coding for trRNA.

**Table 1.** Enrichment of MS-associated genetic variants in lists of T-cell transient transcripts extracted from Michel et al. (21). The whole transcriptome list was split in two sub-lists depending on the transcripts’ half-life: short (<60’) and long (≥60’), respectively. Results are considered significant at p<0.05 and are highlighted in bold.

| List | -log (p-value) | p-value | Odds Ratio | Support | List Size |
| --- | --- | --- | --- | --- | --- |
| Whole transient transcriptome | 8.55 | **$2.80 \times 10^{-9}$** | 1.65 | 241 | 22126 |
| Short half-life transcripts | 7.68 | **$2.06 \times 10^{-8}$** | 1.63 | 209 | 20143 |
| Long half-life transcripts | 1.05 | 0.09 | 1.29 | 35 | 1993 |

When we further dissected the mapping of the ROI colocalization signals, we found a significant excess of intergenic and intron regions (as anticipated), as well as their prevalent distribution away from the transcription start site (TSS; Figure 1A). Notably, when we extended this analysis to GWAS data coming from other multifactorial diseases or traits, dividing immune-mediated and other complex conditions, we found highly comparable profiles (Figure 1B, 1C, Supplementary Table S2), suggesting that the colocalization between MS-associated DNA intervals and intergenic or intronic sequences, plausibly referring to trRNA coding regions, is shared by the genetic architecture of most multifactorial disorders.

**Figure 1.**
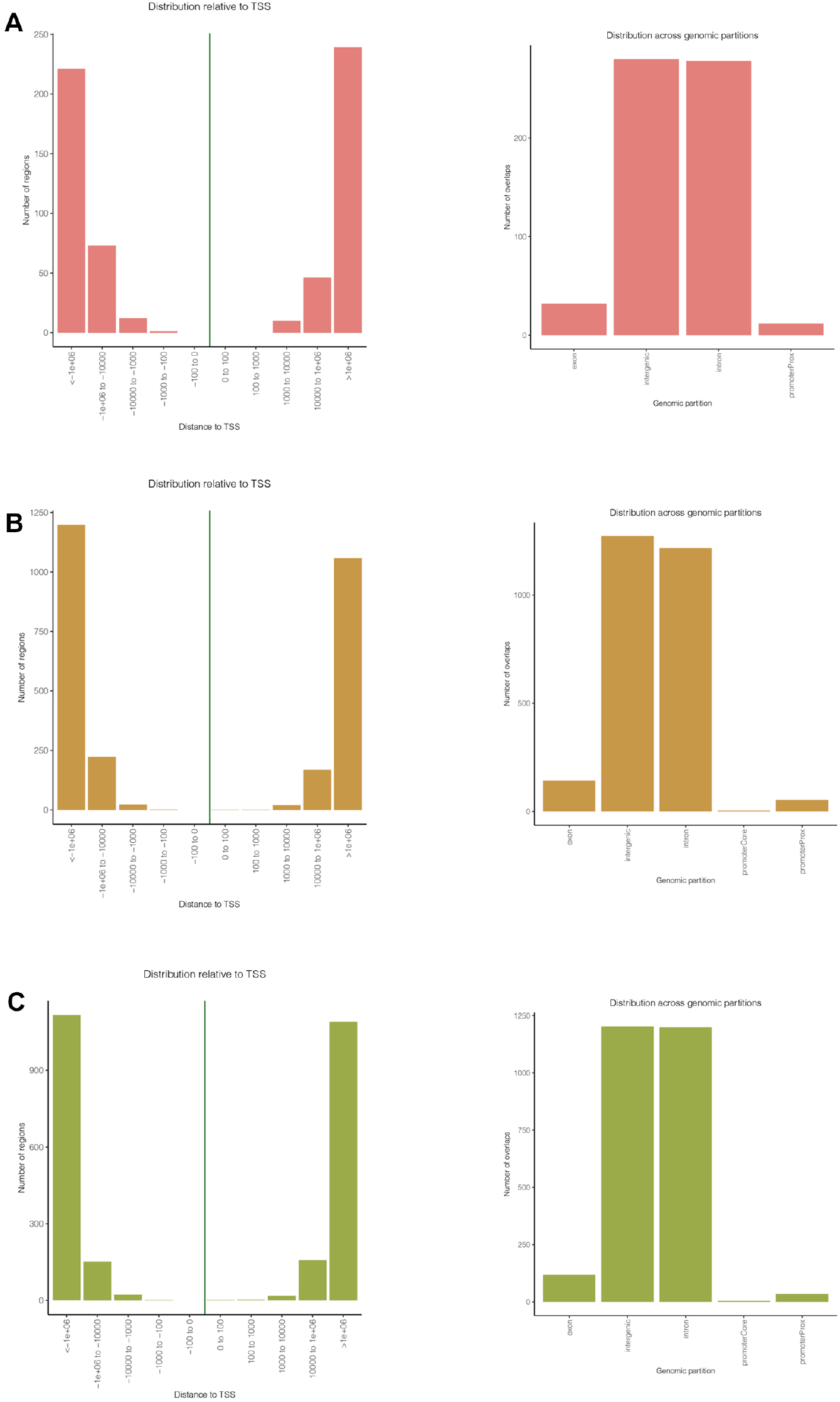
GWAS-associated SNP distribution across genomic partitions and their distance relative to the transcription starting site (TSS). Panel A, Multiple Sclerosis; B, Immune-mediated conditions: Multiple Sclerosis, Rheumatoid Arthritis, Systemic Lupus Erythematosus, Crohn’s Disease, Ulcerative Colitis, Inflammatory Bowel Disease, Celiac Disease, Asthma, Type I Diabetes Mellitus; C, Non-immunological complex conditions: Type II Diabetes Mellitus, Aging, Obesity, Hypertension, Coronary Artery Disease, Bipolar Disorder. Supplementary Table S2 include links to these traits in the GWAS catalog.

To consolidate this result and gain a deeper biological insight, we extended the colocalization analysis matching the ROI with a vast set of databases of regulatory DNA regions, including enhancers and super-enhancers, derived from experiments on diverse tissue types (a total of 4,697,782 DNA regions, plausibly coding for trRNA, were extracted from a wide variety of raw data sources; referenced in Supplementary Table S3). To improve interpretability of the results through ranking, we implemented a harmonic score (HS), based on the Odd Ratio, the −log (p-value), and the support of each match. Statistically significant results came from sets included in SEA, seDB, dbSuper and other single lists of enhancers and non-coding RNAs (Figure 2).

**Figure 2.**
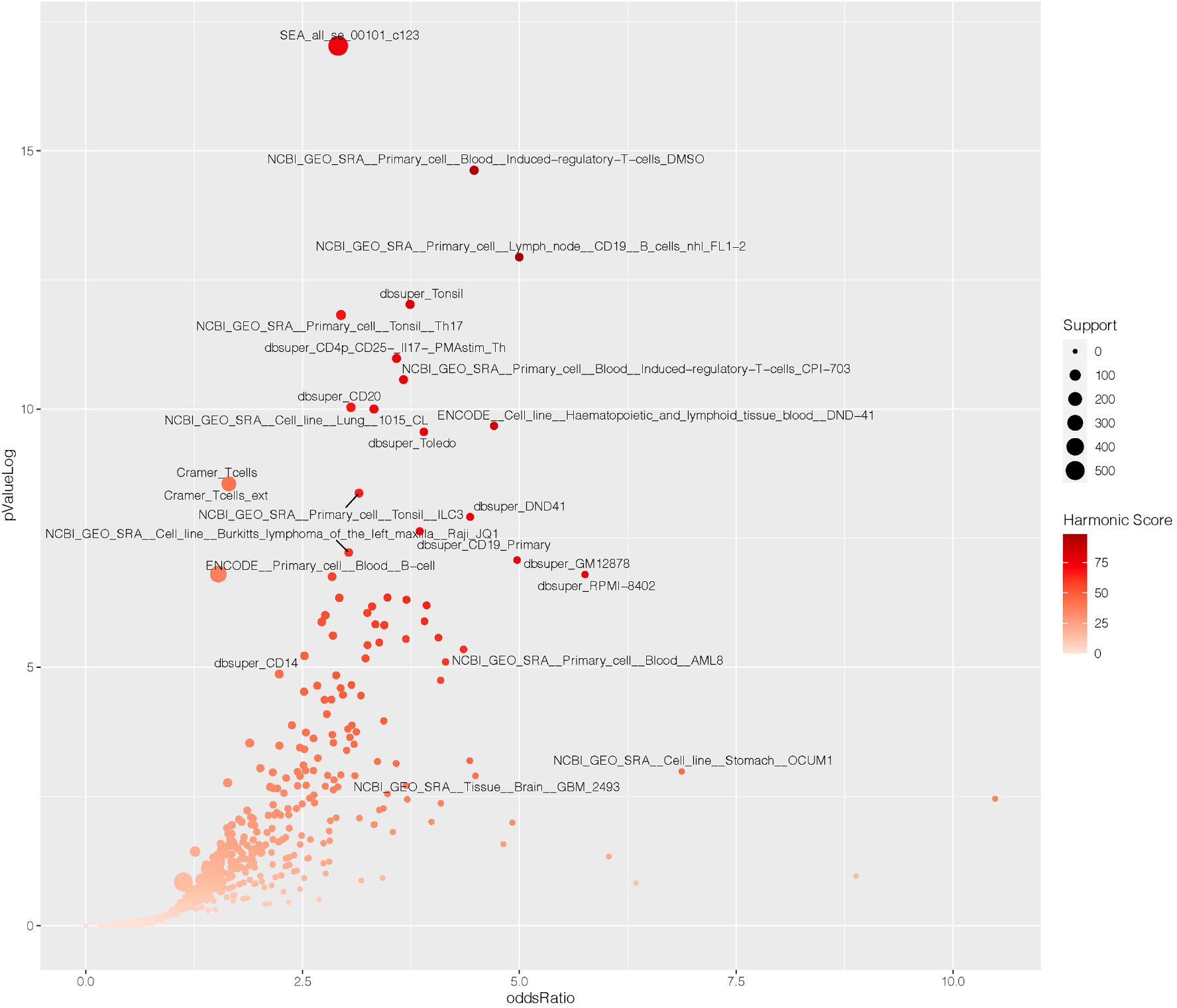
Enrichment of MS-associated SNPs in databases of regulatory elements, sorted by experiment/cell lines. X-axis shows the Odd Ratio, y-axis shows the −log (pValue); dot size is proportional to the support of each match, i.e., the number of hits resulting from the colocalization analysis. Color of each point is related to the Harmonic Score (HS), a comprehensive estimation of the relevance of hits, as derived by merging and balancing the OR, pValue and Support of each match. Thus, prioritized hits are represented by the darker dots that occupy the upper-right area of the chart. Labeled points have HS>40. Labels were arbitrarily designated according to the database of origin and the cell lineage where the enrichment occurred.

Interestingly, we found a strong enrichment of MS-associated genetic variants in cell lines of hematopoietic lineage, including CD19+ and CD20+ B lymphocytes, CD4+ T helper cells, and CD14+ monocytes. Moreover, among the top-scoring hits, we found microglial-specific enhancers, which is in line with recent reports on brain cell type-specific enhancer-promoter interactome activities and the latest GWAS on MS genomic mapping (Consortium, 2019; Nott et al., 2019). On the other hand, non-relevant tissues serving as controls (such as kidney, muscle, glands, etc.) scored low in the ranking, crowding the bottom-left corner of Figure 2.

### Genetic and non-genetic factors for MS etiology converge in genomic regions plausibly coding for the transient transcriptome

Independent studies support the fact that MS GWAS intervals are enriched with DNA binding regions (DBRs) for protein ‘transducers’ mediating non-genetic factors of putative etiologic relevance in MS, such as vitamin D deficiency or EBV latent infection (Harley et al., 2018; Ricigliano et al., 2015). Therefore, we further inquired whether DNA regions plausibly coding for trRNA would share these features (i.e., they colocalize with such DRBs). We set up 4 new ROIs corresponding to the DBRs for VDR, activation-induced cytidine deaminase (AID), EBNA2, and Epstein Barr nuclear antigen 3 (EBNA3C), chosen among viral or host’s nuclear factors potentially associated to MS etiopathogenesis (Bäcker-Koduah et al., 2020; Marcucci & Obeidat, 2020; Sun et al., 2013). The DBRs for each nuclear factor were derived from recent literature (Supplementary Table S4) and matched with the GWAS-derived MS signals to confirm and expand previous results. We found statistically significant results for VDR, EBNA2, and AID for all the SNP position extensions (±50, 100, 200 kb up- and down-stream), while for EBNA3C significant results came out at extension of ±100 and 200 kilobases. This finding suggests that several DBRs can impact on the MS-associated DNA intervals through colocalization (Table 2). Building once again on the work by Michel et al. (Michel et al., 2017), we inquired whether there was a colocalization between genomic regions containing MS-associated variants, DBRs for VDR, EBNA2, EBNA3C, AID, and DNA intervals plausibly coding for trRNA. To this end, we considered the transient transcriptome that proved to be enriched with MS-associated variants (Table 1), and we then matched the corresponding coding regions with the DBRs for the four molecular transducers. For this analysis DBRs for EBNA2 (6,880 regions), EBNA3C (3,835 regions), AID (4,823 regions), and VDR (23,409 regions), represented the ROI, while the ENCODE database of Transcription Factors Binding Sites served as Universe (13,202,334 regions; Figure 3a). We report the results of this analysis in Table 3, which shows the significant colocalization of DNA regions plausibly coding for trRNA with both MS-relevant GWAS signals, and DBR of 3 out of 4 factors active at nuclear level, and potentially associated with MS. The DBR for EBNA3C did not reach statistical significance, though it showed higher values of support for short half-life transcripts.

**Table 2.** Enrichment of MS-GWAS regions (at ±50,100,200 kb range of extension) in lists (number in brackets in the right-most column) of DNA binding sites of human and viral molecular transducers; significant results (p<0.05, corresponding to a −log (p)>1.301) in bold.

| | $\pm 50$ KB | | | | $\pm 100$ KB | | | | $\pm 200$ KB | | | |
| --- | --- | --- | --- | --- | --- | --- | --- | --- | --- | --- | --- | --- |
| | $-\log$<br>(pValue) | Odds Ratio | Support | Harmonic<br>Score | $-\log$<br>(pValue) | Odds Ratio | Support | Harmonic<br>Score | $-\log$<br>(pValue) | Odds Ratio | Support | Harmonic<br>Score |
| **EBNA2**<br>**(6880)** | **10.658** | **1.790** | **158** | **45.544** | **8.616** | **1.509** | **239** | **38.327** | **15.444** | **1.542** | **421** | **41.913** |
| **EBNA3C**<br>**(3335)** | **0.614** | **1.108** | **55** | **11.765** | **1.647** | **1.227** | **109** | **20.956** | **3.448** | **1.294** | **199** | **28.098** |
| **AID**<br>**(4823)** | **4.963** | **1.596** | **99** | **35.793** | **3.890** | **1.374** | **153** | **30.259** | **13.924** | **1.619** | **309** | **43.308** |
| **VDR**<br>**(23409)** | **19.348** | **1.575** | **474** | **43.564** | **19.181** | **1.422** | **767** | **39.635** | **32.090** | **1.424** | **1329** | **40.872** |

**Figure 3.**
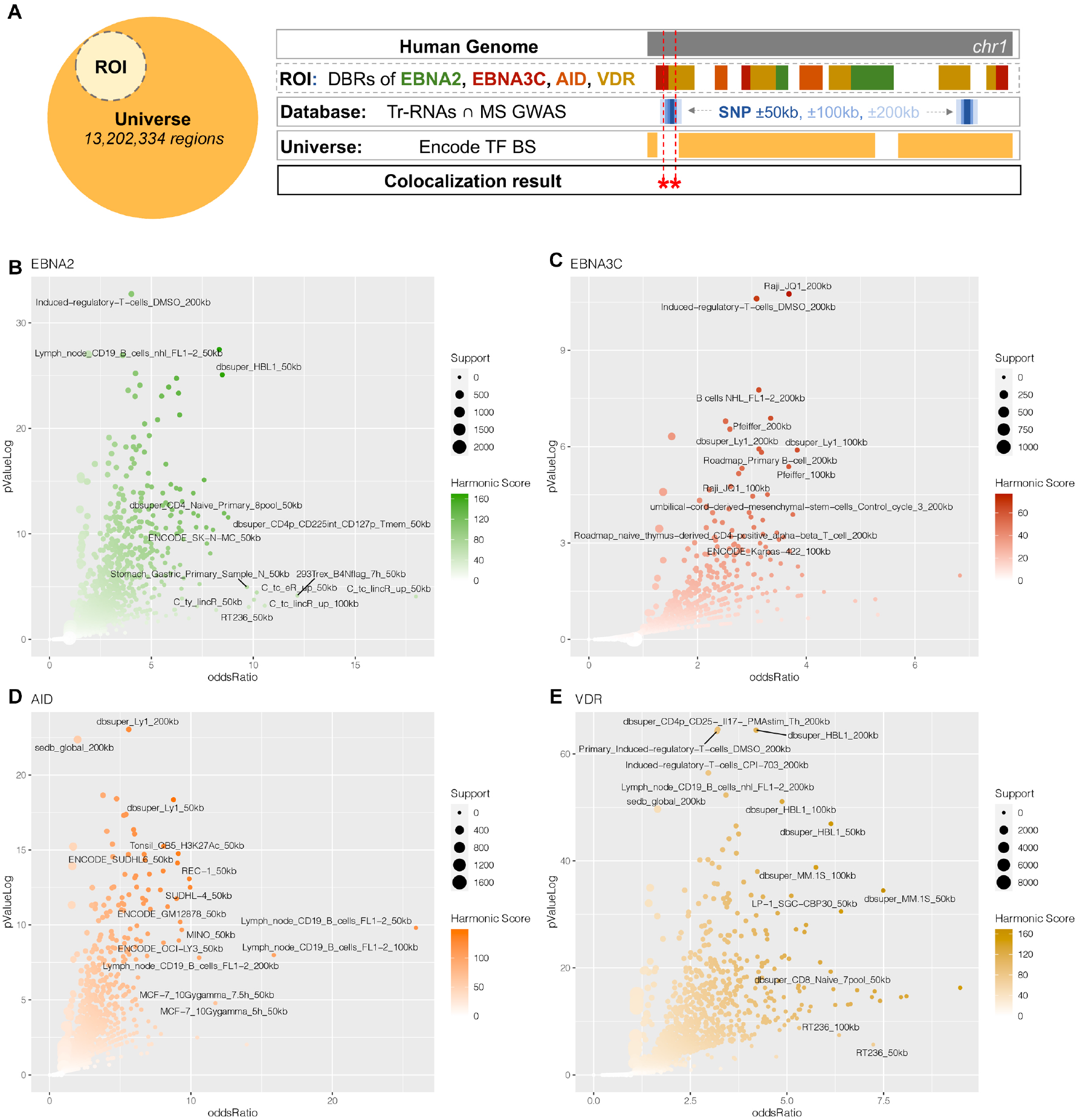
Colocalization analysis of DBRs for human and viral molecular transducers, MS-associated SNPs and DNA regulatory regions derived from databases. (*A),*Schematic representation of the colocalization analysis. (*ROI: Region of interest; DBR: DNA Binding Regions; ENCODE TFBS: Transcription Factor Binding Site).* The figure shows the tracks we considered for the colocalization analyses. In brief, the ROI included the DBRs of MS-related viral and human transducers and was matched with MS-associated SNPs extended by 50, 100, and 200 kilobases that colocalize with regions plausibly coding for trRNAs (Database). As a control (Universe), we took from ENCODE the entire list of transcription factors binding sites. Results were considered significant if a colocalization was found across ROI and a Databases element without occurring in the Universe as a statistically significant match*. (B-E*), Colocalization results of EBNA2, EBNA3C, AID, VDR. The charts display results of all matches, i.e, with MS-associated SNPs and their extension at ±50, 100, 200 kb. X-axis shows the Odd Ratio, y-axis shows the log (pValue). Dot size is proportional to the support of each match, i.e., the number of hits resulting from each colocalization analysis. The color of each dot is related to the Harmonic Score (HS); labeled points have HS>40. Labels were arbitrarily designated according to the database of origin and the cell lineage where the enrichment occurred.

**Table 3.** Colocalization of human and viral transducer DBRs and MS-GWAS positions (at ±50,100,200 kb range of extension) in DNA regions coding for transient transcripts; significant results (p<0.05, corresponding to a −log (p)>1.301) in bold. The transcript half-life is considered short if <60’ and long if ≥60’, respectively.

| | | $\pm 50$ KB | | | | $\pm 100$ KB | | | | $\pm 200$ KB | | | |
| --- | --- | --- | --- | --- | --- | --- | --- | --- | --- | --- | --- | --- | --- |
| | | $-\log$<br>(pValue) | Odds<br>Ratio | Support | Harmonic<br>Score | $-\log$<br>(pValue) | Odds<br>Ratio | Support | Harmonic<br>Score | $-\log$<br>(pValue) | Odds<br>Ratio | Support | Harmonic<br>Score |
| **EBNA2** | *Long half-life* | 0.023 | 0.478 | 3 | 0.644 | 0.062 | 0.717 | 8 | 1.708 | **1.879** | **1.531** | **33** | **24.679** |
|  | *Short half-life* | **6.163** | **1.920** | **69** | **43.011** | **3.241** | **1.433** | **95** | **29.496** | **8.945** | **1.610** | **189** | **40.642** |
| **EBNA3C** | *Long half-life* | 0.064 | 0.572 | 2 | 1.669 | 0.006 | 0.321 | 2 | 0.185 | 0.182 | 0.914 | 11 | 4.500 |
|  | *Short half-life* | 0.070 | 0.794 | 16 | 1.923 | 0.023 | 0.752 | 28 | 0.661 | 0.066 | 0.875 | 58 | 1.841 |
| **AID** | *Long half-life* | 0.089 | 0.682 | 3 | 2.303 | 0.283 | 1.024 | 8 | 6.477 | 0.051 | 0.726 | 11 | 1.432 |
|  | *Short half-life* | **1.769** | **1.465** | **37** | **23.531** | **1.346** | **1.267** | **59** | **19.367** | **3.954** | **1.442** | **119** | **31.416** |
| **VDR** | *Long half-life* | **1.737** | **1.502** | **32** | **23.571** | 0.845 | 1.187 | 45 | 14.646 | **2.315** | **1.322** | **97** | **25.031** |
|  | *Short half-life* | **2.221** | **1.239** | **152** | **23.734** | **2.336** | **1.181** | **267** | **23.460** | **11.478** | **1.367** | **548** | **36.561** |

To review and confirm previous colocalizations, we considered the genomic regions resulting from the above reported match between the MS-associated GWAS intervals and the databases of regulatory DNA regions, containing enhancers and super-enhancers, plausibly enriched in trRNA-coding sequences (see results in Figure 2 and the online data resource). We therefore matched these DNA regions with the DBR for VDR, EBNA2, EBNA3C and AID, finding significant enrichments that allow to contextualize and prioritize genomic positions, cell/tissue identity or cell status associated to MS. Considering the harmonic score obtained from these colocalization analyses, the top hits in EBNA2, EBNA3C, and AID involved lymphoid (CD19+ B cell lines and lymphomas; T regulatory cells; tonsils) and monocyte-macrophage lineages (peripheral macrophages; dendritic cells) from experiments included in the ENCODE, dbsuper, roadmapEpigenomics databases (Figure3B-E, see also Supplementary Table S5). Even though immune cells prevailed also in VDR top hits, a less stringent polarization was seen, somehow reflecting the wide-spreading actions of this transducer in human biology. However, with a more stringent cutoff of Harmonic Score>40 that selects the most significant hits (Figure Supplement 1), a core subset of MS-relevant cell lineages, shared across all four examined transducers, became evident (Supplementary Table S6 and the online data resource).

### A data resource for future research on transcriptional regulation in MS

A public web interface for browsing the results of our colocalization analysis is freely available at www.mscoloc.com. This is a comprehensive genomic atlas disentangling specific aspects of MS gene-environment interactions to support further research on transcriptional regulation in MS. It includes the whole list of results derived from ROI, DBRs and database matches (Figure 4a) across all performed experiments that yielded significant results. The user can navigate across the results and perform tailored queries searching and filtering for a variety of parameters, including MS-associated variant, DBR, experimental cell type, other match details (see Figure 4b for all available search and filter modalities). Moreover, personalized HS, p-value, support and Odd Ratio threshold can easily be set to screen results, that are readily displayed in tabular format. To provide an example, we select *“AID, EBNA2, EBNA3C, VDR”* in the ‘*Matched DBR region (s)*’ panel and obtain the list of MS-associated SNPs targeted by all four transducers (Figure 4b-c). Through this approach we searched for MS-associated regions shared by the DBRs analyzed, and we were able to prioritize 275 genomic regions (almost half of the MS-associated GWAS SNPs) capable of binding at least 2 molecular transducers. These regions are ‘hotspots’ of interactions between genetic and nongenetic modifier of MS risk/protection: all four proteins (VDR, AID, EBNA2, EBNA3C) proved to target 24 regions, 3 of them 115 regions, and 2 of them 136 regions. A detailed legend and more example queries may be found on the online data resource website.

**Figure 4.**
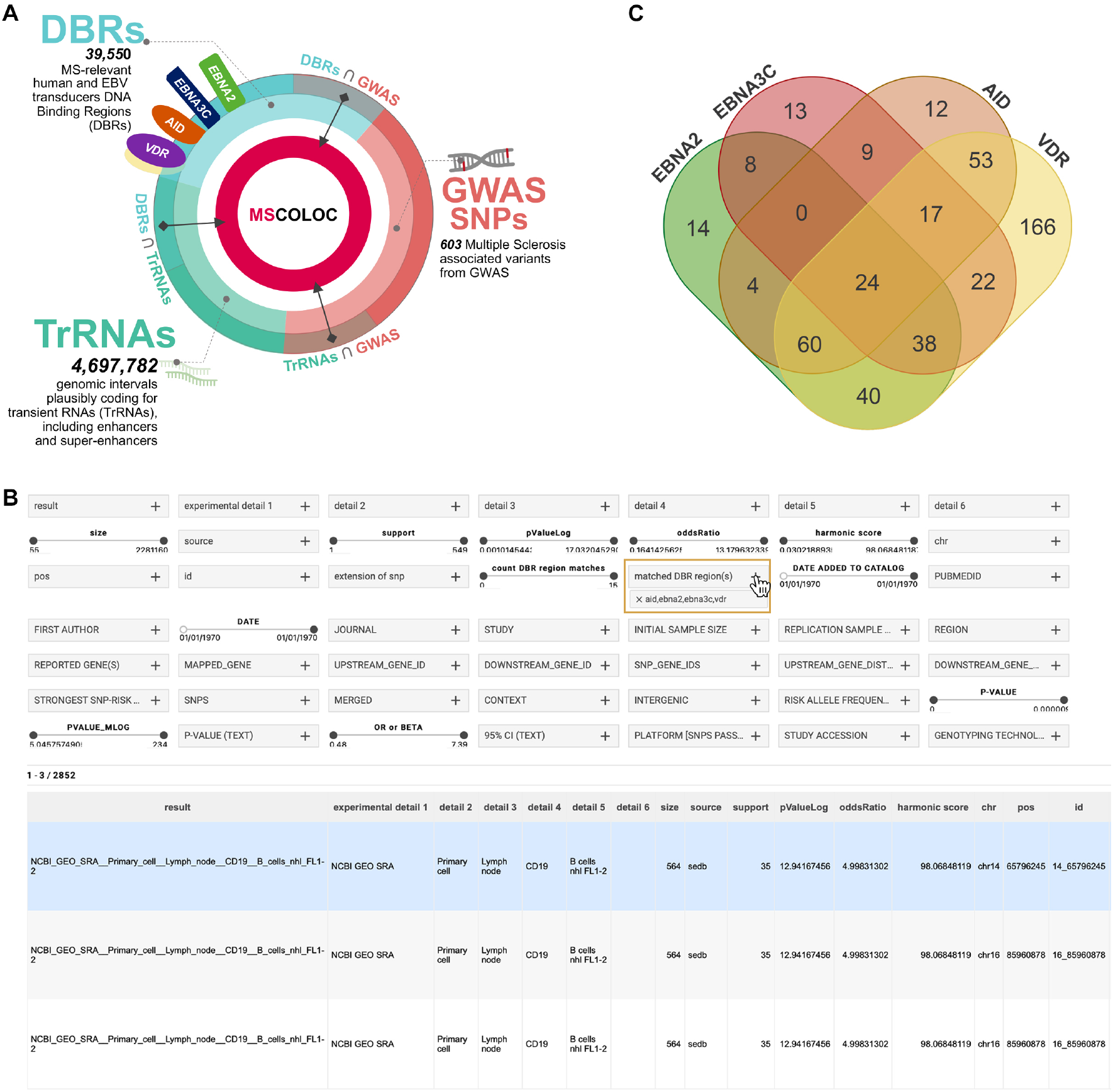
A comprehensive genomic atlas on gene-environment interactions regulating transcription in MS. (*A),* Searchable results at mscoloc.com derive from the matches of GWAS MS regions, DNA binding regions of selected genomic transducers, and more than 4 million of regions annotated as plausible transient RNAs. (*B),* The user interface includes text panels and range sliders allowing extremely personalized queries, that combine statistical significance level (including Odd Ratio, pValue, support, and Harmonic Score), study source, SNP or reported gene, and so on. Filtered results are shown as tables ranked by HS, that can be saved, printed or shared through URL. In the example, the cursor selects ‘*AID, EBNA2, EBNA3C, VDR’* in the ‘*matched DBR region (s)’* panel looking for MS-associated SNPs (from the ROI, Supplementary Table S1) and their extensions at ±50, 100, 200 kb that colocalized within DNA binding regions of the molecular transducers. The top hit represents the colocalization of the DBRs, a super-enhancer region derived from experiments on CD19+B cells included in *sedb,* and the rs8007846 MS-associated SNP on chromosome 14. (*C),* The Venn diagram shows the number of non-redundant MS-associated SNPs derived from the query: for each transducer, SNPs are considered only once if present in more than one match. Intersections show the numbers of regions colocalizing with DBRs of multiple transducers. For instance: 8 regions colocalize with both EBNA2 and EBNA3C DBRs, but not with AID nor VDR DBRs; 24 regions colocalize with all four DBRs, and could be identified as regulatory “hotspots” in MS.

## Discussion

Our study supports the hypothesis that investigations on the transient transcriptome may contribute to clarify how the GWAS signals affect the etiopathogenesis of MS and possibly of other complex disorders. Specifically, we show that genomic regions coding for the transient transcriptome recently described in T cells (Michel et al., 2017), are significantly enriched for both MS-associated GWAS variants, as well as for DNA binding sites for protein ‘transducers’ of non-genetic signals, chosen among those plausibly associated to MS. The colocalization of GWAS intervals and some DNA-binding factors involved in MS etiology has already been reported (Harley et al., 2018; Mechelli et al., 2015; Ricigliano et al., 2015), and here we reinforce this premise further suggesting a model in which trRNA-coding regions are hotspots of convergence between genetic ad non-genetic factors of risk/protection for MS. Our analysis showed that these hotspots are shared by two or more of the chosen transducers, indicating possible additive pathogenic effects or a multi-hits model to reach the threshold for MS development (see Figure 4 and Supplementary Table S6).

In homeostatic conditions, it can be hypothesized that DNA sequences coding for trRNA are composed of regulatory regions where genetic variability and non-genetic signals interact to finely regulate the gene expression according to cell identity, developmental or adaptive states, and time-dependent stimuli. As a matter of fact, the sequence variability of these regions and the strict time-dependence of their transcription could be instrumental to adaptive features; however, these same features make these regions susceptible to become dysfunctional or to be the targets of pathogenic interaction. In some instances, these detrimental interactions come from outside the cell, such as in the case of EBV interference with host transcription (Mechelli R., 2021, Accepted; Park et al., 2020), and the pathogenic consequences of vitamin D deficiency; in other cases, the dysfunction develops within the cell, such as the tumorigenic activity of AID in B cells (Meng et al., 2014; Qian et al., 2014).

The mapping of transient transcripts by TT-seq approach fits very well with our results obtained from GWAS data for MS and other multifactorial conditions, showing a significant excess of intergenic and intronic regions (coding for eRNA, sincRNA, and asRNA), and having a distribution in DNA intervals mostly far off from transcription start sites (TSS; see Figure 1). This is in agreement with recent evidence of regulatory DNA region markers which contain genetic variants for complex disease or traits; indeed, a systematic framework of common coordinates for these markers showed that about half of them lie within introns and most are located away from the TSS (Meuleman et al., 2020).

To further support the relationship between trRNA and transcription of regulatory DNA regions, we matched a large dataset of enhancers and super-enhancers with MS-GWAS signals and DBR for VDR, EBNA2, EBNA3C and AID. The significant enrichment in cell lines and cell status coming from the hematopoietic lineages and the CNS-specific cell subsets corroborates data coming from recent reports showing the relevance of contextualizing and prioritizing the role of MS-associated GWAS signals (Consortium, 2019; Factor et al., 2020; Nott et al., 2019; Orrù et al., 2020). Our analysis supports the pivotal regulatory role of enhancer transcription (i.e., a main component of transient transcriptome) that was recently reported as not dispensable for gene expression at the immunoglobulin locus and for antibody class switch recombination (Fitz et al., 2020), though more research is needed to unravel such topic at a finer grain.

Reports on the dynamics of time-course data are a recent area of focus within the analysis of gene expression, specifically in immune cells. Although current studies use methods that investigate time points related to the stable transcriptome (RNA-seq performed with time spans of hours), they clearly show that gene expression dynamics may influence allele specificity, regulatory programs that seem to depend on autoimmune disease-associated loci, and different transcriptional profiles based on cell status after stimulation (Gutierrez-Arcelus et al., 2020). A recent work showed that an *IL2ra* enhancer, which harbors autoimmunity risk variants and was one of the first MS-associated loci from GWAS, has no impact on the gene level expression, but rather affects gene activation by delaying transcription in response to extracellular stimuli (Simeonov et al., 2017). The importance of the timing in the gene expression control emerges also from several studies implicating enhancers and super-enhancers in the process of phase separation and formation of condensates. In this context, the transcriptional apparatus steps-up to drive robust genic responses (. The overall process seems to be highly dynamic, with time spans of seconds or minutes, and hence compatible with the temporal features of the transient transcriptome, which could somehow act upstream for the formation of these phase-separated condensates.

We suggest that studies on transient transcriptomes may integrate previous RNA-seq data in accounting for the interplay between genetic variability and non-genetic etiologic factors leading to MS development. Components of a more-complex-than-anticipated regulation of gene expression could include transcriptional noise, transitory time-courses, erratic dynamics, and highly flexibility of some DNA regions, possibly oscillating between bistable states of enhancer and silencer (Halfon, 2020). The availability of tools to map trRNA could further contribute to the development of studies on immune cells isolated from patients and matched controls, aimed at dissecting key aspects of the complex transcriptional response in MS. Our analysis provides a platform for future studies on transient transcriptome, which we support by making our data resource available at www.mscoloc.com. Finally, new gene regulatory models may emerge from this approach in order to better evaluate the meaning of GWAS in complex traits and the impact of the enhancer transcription (Fitz et al., 2020), which was recently reported as an ancient and conserved, yet flexible, genomic regulatory syntax (Wong et al., 2020).

## Materials and Methods

### Data pipeline

Analyses were performed in Python and R. A data freeze was applied on 3/1/2020. All GWAS data was gathered from the GWAS Catalog through its REST API (Buniello et al., 2019); about 1.5% of this data was filtered out as part of a QC process aimed at homogenizing legacy and more recent data. The MS GWAS regions were extracted from the overall GWAS Catalog data filtering by trait EFO_0003885. All Transcription Factor Binding Site regions (TFBS) were obtained from the ENCODE portal (Sloan et al., 2016). All data was organized in various databases and data pipelines as detailed below. A modular and parallel data pipeline was created to: (i) readily generate and evaluate all experiments in the paper, (ii) manage and organize all data coming from various region collections (42,075 ROI regions; 4,697,782 regions plausibly coding for trRNAs; 13,309,757 Universe regions), multiple ROIs (MS GWAS, EBNA2, EBNA3C, VDR, AID, etc.), databases of vast background regions as they were populated with the data obtained from GWAS Catalog, ENCODE, and other raw data sources, (iii) provide overlaps and intersection among various data elements, annotate them with the original MS GWAS loci that generated the signal, and (iv) generate the overarching data resource available at www.mscoloc.com.

### Statistical analysis

For SNP overlaps and region colocalization, we used LOLA (Sheffield & Bock, 2016) and Fisher’s exact test with False Discovery Rate (Benjamini-Hochberg) to control for multiple testing. Resulting −log (p-value), support, and Odds Ratio (OR) were combined into a single score inspired by the harmonic mean (Wilson, 2019) and multi-objective optimization (Umeton R., 2011) with the formula below, where the spacing parameter *k_p_* was set to 10.0 and we consider all three contributors equally, setting therefore weights *w_i_* to 1.0. Statistical significance was taken at p<0.05.

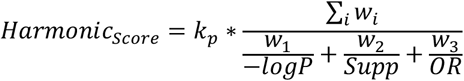

## Data availability

All generated data and results are made available at the website www.mscoloc.com.

## Author Contributions

RU, GB, RB, RM, MS, and GR conceived and planned the analysis. RR, VR, EM, CR, SR, and MCB guided data engineering and database generation from raw data. RU, RPU and GB developed the data resource. RU, GB, RPU, GR and MS wrote the manuscript. RU, GB, RB, and RM created all table and figures. RB, RM, MS and GR supervised the project. All the authors, including SR, MCB, RR, VR, EM and CR, contributed to fortnight discussion for data interpretation and new analysis planning.

All the authors revised and approved the manuscript.

## Competing Interest Statement

The authors have no conflict of interest related to this analysis.

## Acknowledgments

This work was supported by “Progetti Grandi Ateneo” 2020, Sapienza University of Rome. MS and GR are supported by CENTERS, a Special Project of, and financed by, FISM - Fondazione Italiana Sclerosi Multipla.

## Supplementary material

### Supplementary Tables

**S1**. MS-associated genomic positions from GWAS catalog after QC process filtering, used as Region of Interest (ROI) for the analysis.

**S2**. GWAS Catalog References for diseases considered in Figure 1.

**S3**. Sources of DNA regions plausibly coding for trRNAs with references.

**S4**. Sources of DNA Binding Regions (DBRs) of considered viral and human transducers with references.

**S5**. Top 10 results of Colocalization analysis.

**S6**. Cell types for which the colocalization analysis hits reported a harmonic score >40 in all transducers (EBNA2, EBNA3C, AID, VDR).

**Figure supplement 1:** Harmonic Score threshold defining the top colocalization hits.

## Notes

https://www.mscoloc.com

### Competing Interest Statement

The authors have declared no competing interest.

### Summary of Updates

- Typo fixed

## References

Bordi, I., Ricigliano, V. A., Umeton, R., Ristori, G., Grassi, F., Crisanti, A., Sutera, A., & Salvetti, M. (2014). Noise in multiple sclerosis: unwanted and necessary. Ann Clin Transl Neurol, 1 (7), 502–511. https://doi.org/10.1002/acn3.72

Bordi, I., Umeton, R., Ricigliano, V. A., Annibali, V., Mechelli, R., Ristori, G., Grassi, F., Salvetti, M., & Sutera, A. (2013). A mechanistic, stochastic model helps understand multiple sclerosis course and pathogenesis. Int J Genomics, 2013, 910321. https://doi.org/10.1155/2013/910321

Bose, D. A., Donahue, G., Reinberg, D., Shiekhattar, R., Bonasio, R., & Berger, S. L. (2017). RNA Binding to CBP Stimulates Histone Acetylation and Transcription. Cell, 168 (1-2), 135–149.e122. https://doi.org/10.1016/j.cell.2016.12.020

Buniello, A., MacArthur, J. A. L., Cerezo, M., Harris, L. W., Hayhurst, J., Malangone, C., McMahon, A., Morales, J., Mountjoy, E., Sollis, E., Suveges, D., Vrousgou, O., Whetzel, P. L., Amode, R., Guillen, J. A., Riat, H. S., Trevanion, S. J., Hall, P., Junkins, H., Flicek, P., Burdett, T., Hindorff, L. A., Cunningham, F., & Parkinson, H. (2019). The NHGRI-EBI GWAS Catalog of published genome-wide association studies, targeted arrays and summary statistics 2019. Nucleic Acids Res, 47 (D1), D1005–D1012. https://doi.org/10.1093/nar/gky1120

Bäcker-Koduah, P., Bellmann-Strobl, J., Scheel, M., Wuerfel, J., Wernecke, K. D., Dörr, J., Brandt, A. U., & Paul, F. (2020). Vitamin D and Disease Severity in Multiple Sclerosis-Baseline Data From the Randomized Controlled Trial (EVIDIMS). Front Neurol, 11, 129. https://doi.org/10.3389/fneur.2020.00129

Calderon, D., Nguyen, M. L. T., Mezger, A., Kathiria, A., Müller, F., Nguyen, V., Lescano, N., Wu, B., Trombetta, J., Ribado, J. V., Knowles, D. A., Gao, Z., Blaeschke, F., Parent, A. V., Burt, T. D., Anderson, M. S., Criswell, L. A., Greenleaf, W. J., Marson, A., & Pritchard, J. K. (2019). Landscape of stimulation-responsive chromatin across diverse human immune cells. Nat Genet, 51 (10), 1494–1505. https://doi.org/10.1038/s41588-019-0505-9

, I. M. S. G. C. E. a., & Consortium, I. M. S. G. (2020). Low-Frequency and Rare-Coding Variation Contributes to Multiple Sclerosis Risk. Cell, 180 (2), 403. https://doi.org/10.1016/j.cell.2020.01.002

Consortium, G. (2020). The GTEx Consortium atlas of genetic regulatory effects across human tissues. Science, 369 (6509), 1318–1330. https://doi.org/10.1126/science.aaz1776

Consortium, I. M. S. G. (2019). Multiple sclerosis genomic map implicates peripheral immune cells and microglia in susceptibility. Science, 365 (6460). https://doi.org/10.1126/science.aav7188

Ernst, J., Kheradpour, P., Mikkelsen, T. S., Shoresh, N., Ward, L. D., Epstein, C. B., Zhang, X., Wang, L., Issner, R., Coyne, M., Ku, M., Durham, T., Kellis, M., & Bernstein, B. E. (2011). Mapping and analysis of chromatin state dynamics in nine human cell types. Nature, 473 (7345), 43–49. https://doi.org/10.1038/nature09906

Factor, D. C., Barbeau, A. M., Allan, K. C., Hu, L. R., Madhavan, M., Hoang, A. T., Hazel, K. E. A., Hall, P. A., Nisraiyya, S., Najm, F. J., Miller, T. E., Nevin, Z. S., Karl, R. T., Lima, B. R., Song, Y., Sibert, A. G., Dhillon, G. K., Volsko, C., Bartels, C. F., Adams, D. J., Dutta, R., Gallagher, M. D., Phu, W., Kozlenkov, A., Dracheva, S., Scacheri, P. C., Tesar, P. J., & Corradin, O. (2020). Cell Type-Specific Intralocus Interactions Reveal Oligodendrocyte Mechanisms in MS. Cell, 181 (2), 382–395.e321. https://doi.org/10.1016/j.cell.2020.03.002

Fagnani, C., Neale, M. C., Nisticò, L., Stazi, M. A., Ricigliano, V. A., Buscarinu, M. C., Salvetti, M., & Ristori, G. (2015). Twin studies in multiple sclerosis: A meta-estimation of heritability and environmentality. Mult Scler, 21 (11), 1404–1413. https://doi.org/10.1177/1352458514564492

Farh, K. K., Marson, A., Zhu, J., Kleinewietfeld, M., Housley, W. J., Beik, S., Shoresh, N., Whitton, H., Ryan, R. J., Shishkin, A. A., Hatan, M., Carrasco-Alfonso, M. J., Mayer, D., Luckey, C. J., Patsopoulos, N. A., De Jager, P. L., Kuchroo, V. K., Epstein, C. B., Daly, M. J., Hafler, D. A., & Bernstein, B. E. (2015). Genetic and epigenetic fine mapping of causal autoimmune disease variants. Nature, 518 (7539), 337–343. https://doi.org/10.1038/nature13835

Fitz, J., Neumann, T., Steininger, M., Wiedemann, E. M., Garcia, A. C., Athanasiadis, A., Schoeberl, U. E., & Pavri, R. (2020). Spt5-mediated enhancer transcription directly couples enhancer activation with physical promoter interaction. Nat Genet, 52 (5), 505–515. https://doi.org/10.1038/s41588-020-0605-6

Gusev, A., Lee, S. H., Trynka, G., Finucane, H., Vilhjálmsson, B. J., Xu, H., Zang, C., Ripke, S., Bulik-Sullivan, B., Stahl, E., Kähler, A. K., Hultman, C. M., Purcell, S. M., McCarroll, S. A., Daly, M., Pasaniuc, B., Sullivan, P. F., Neale, B. M., Wray, N. R., Raychaudhuri, S., Price, A. L., Consortium, S. W. G. o. t. P. G., & Consortium, S.-S. (2014). Partitioning heritability of regulatory and cell-type-specific variants across 11 common diseases. Am J Hum Genet, 95 (5), 535–552. https://doi.org/10.1016/j.ajhg.2014.10.004

Gutierrez-Arcelus, M., Baglaenko, Y., Arora, J., Hannes, S., Luo, Y., Amariuta, T., Teslovich, N., Rao, D. A., Ermann, J., Jonsson, A. H., Navarrete, C., Rich, S. S., Taylor, K. D., Rotter, J. I., Gregersen, P. K., Esko, T., Brenner, M. B., Raychaudhuri, S., & Consortium, N. T.-O. f. P. M. T. (2020). Allele-specific expression changes dynamically during T cell activation in HLA and other autoimmune loci. Nat Genet, 52 (3), 247–253. https://doi.org/10.1038/s41588-020-0579-4

Halfon, M. S. (2020). Silencers, Enhancers, and the Multifunctional Regulatory Genome. Trends Genet, 36 (3), 149–151. https://doi.org/10.1016/j.tig.2019.12.005

Harley, J. B., Chen, X., Pujato, M., Miller, D., Maddox, A., Forney, C., Magnusen, A. F., Lynch, A., Chetal, K., Yukawa, M., Barski, A., Salomonis, N., Kaufman, K. M., Kottyan, L. C., & Weirauch, M. T. (2018). Transcription factors operate across disease loci, with EBNA2 implicated in autoimmunity. Nat Genet, 50 (5), 699–707. https://doi.org/10.1038/s41588-018-0102-3

Isoda, T., Moore, A. J., He, Z., Chandra, V., Aida, M., Denholtz, M., Piet van Hamburg, J., Fisch, K. M., Chang, A. N., Fahl, S. P., Wiest, D. L., & Murre, C. (2017). Non-coding Transcription Instructs Chromatin Folding and Compartmentalization to Dictate Enhancer-Promoter Communication and T Cell Fate. Cell, 171 (1), 103–119.e118. https://doi.org/10.1016/j.cell.2017.09.001

Larsson, A. J. M., Johnsson, P., Hagemann-Jensen, M., Hartmanis, L., Faridani, O. R., Reinius, B., Segerstolpe, Å., Rivera, C. M., Ren, B., & Sandberg, R. (2019). Genomic encoding of transcriptional burst kinetics. Nature, 565 (7738), 251–254. https://doi.org/10.1038/s41586-018-0836-1

Marcucci, S. B., & Obeidat, A. Z. (2020). EBNA1, EBNA2, and EBNA3 link Epstein-Barr virus and hypovitaminosis D in multiple sclerosis pathogenesis. J Neuroimmunol, 339, 577116. https://doi.org/10.1016/j.jneuroim.2019.577116

Maurano, M. T., Humbert, R., Rynes, E., Thurman, R. E., Haugen, E., Wang, H., Reynolds, A. P., Sandstrom, R., Qu, H., Brody, J., Shafer, A., Neri, F., Lee, K., Kutyavin, T., Stehling-Sun, S., Johnson, A. K., Canfield, T. K., Giste, E., Diegel, M., Bates, D., Hansen, R. S., Neph, S., Sabo, P. J., Heimfeld, S., Raubitschek, A., Ziegler, S., Cotsapas, C., Sotoodehnia, N., Glass, I., Sunyaev, S. R., Kaul, R., & Stamatoyannopoulos, J. A. (2012). Systematic localization of common disease-associated variation in regulatory DNA. Science, 337 (6099), 1190–1195. https://doi.org/10.1126/science.1222794

Mechelli, R., Manzari, C., Policano, C., Annese, A., Picardi, E., Umeton, R., Fornasiero, A., D’Erchia, A. M., Buscarinu, M. C., Agliardi, C., Annibali, V., Serafini, B., Rosicarelli, B., Romano, S., Angelini, D. F., Ricigliano, V. A., Buttari, F., Battistini, L., Centonze, D., Guerini, F. R., D’Alfonso, S., Pesole, G., Salvetti, M., & Ristori, G. (2015). Epstein-Barr virus genetic variants are associated with multiple sclerosis. Neurology, 84 (13), 1362–1368. https://doi.org/10.1212/WNL.0000000000001420

Mechelli R., R. C., Reniè R., Manfrè G., Buscarinu M.C., Romano S., Marrone A., Bigi R., Bellucci G., Ballerini C., Angeloni B., Rinaldi V., Salvetti M., Ristori G. (2021, Accepted). Viruses and neuroinflammation in multiple sclerosis. In: Neuroimmunology and Neuroinflammation.

Meng, F. L., Du, Z., Federation, A., Hu, J., Wang, Q., Kieffer-Kwon, K. R., Meyers, R. M., Amor, C., Wasserman, C. R., Neuberg, D., Casellas, R., Nussenzweig, M. C., Bradner, J. E., Liu, X. S., & Alt, F. W. (2014). Convergent transcription at intragenic super-enhancers targets AID-initiated genomic instability. Cell, 159 (7), 1538–1548. https://doi.org/10.1016/j.cell.2014.11.014

Meuleman, W., Muratov, A., Rynes, E., Halow, J., Lee, K., Bates, D., Diegel, M., Dunn, D., Neri, F., Teodosiadis, A., Reynolds, A., Haugen, E., Nelson, J., Johnson, A., Frerker, M., Buckley, M., Sandstrom, R., Vierstra, J., Kaul, R., & Stamatoyannopoulos, J. (2020). Index and biological spectrum of human DNase I hypersensitive sites. Nature, 584 (7820), 244–251. https://doi.org/10.1038/s41586-020-2559-3

Michel, M., Demel, C., Zacher, B., Schwalb, B., Krebs, S., Blum, H., Gagneur, J., & Cramer, P. (2017). TT-seq captures enhancer landscapes immediately after T-cell stimulation. Mol Syst Biol, 13 (3), 920. https://doi.org/10.15252/msb.20167507

Mumbach, M. R., Satpathy, A. T., Boyle, E. A., Dai, C., Gowen, B. G., Cho, S. W., Nguyen, M. L., Rubin, A. J., Granja, J. M., Kazane, K. R., Wei, Y., Nguyen, T., Greenside, P. G., Corces, M. R., Tycko, J., Simeonov, D. R., Suliman, N., Li, R., Xu, J., Flynn, R. A., Kundaje, A., Khavari, P. A., Marson, A., Corn, J. E., Quertermous, T., Greenleaf, W. J., & Chang, H. Y. (2017). Enhancer connectome in primary human cells identifies target genes of disease-associated DNA elements. Nat Genet, 49 (11), 1602–1612. https://doi.org/10.1038/ng.3963

Natoli, G., & Andrau, J. C. (2012). Noncoding transcription at enhancers: general principles and functional models. Annu Rev Genet, 46, 1–19. https://doi.org/10.1146/annurev-genet-110711-155459

Nott, A., Holtman, I. R., Coufal, N. G., Schlachetzki, J. C. M., Yu, M., Hu, R., Han, C. Z., Pena, M., Xiao, J., Wu, Y., Keulen, Z., Pasillas, M. P., O’Connor, C., Nickl, C. K., Schafer, S. T., Shen, Z., Rissman, R. A., Brewer, J. B., Gosselin, D., Gonda, D. D., Levy, M. L., Rosenfeld, M. G., McVicker, G., Gage, F. H., Ren, B., & Glass, C. K. (2019). Brain cell type-specific enhancer-promoter interactome maps and disease. Science, 366 (6469), 1134–1139. https://doi.org/10.1126/science.aay0793

O’Donoghue, G. P., Bugaj, L. J., Anderson, W., Daniels, K. G., Rawlings, D. J., & Lim, W. A. (2021). T cells selectively filter oscillatory signals on the minutes timescale. Proc Natl Acad Sci U S A, 118 (9). https://doi.org/10.1073/pnas.2019285118

Ohkura, N., Yasumizu, Y., Kitagawa, Y., Tanaka, A., Nakamura, Y., Motooka, D., Nakamura, S., Okada, Y., & Sakaguchi, S. (2020). Regulatory T Cell-Specific Epigenomic Region Variants Are a Key Determinant of Susceptibility to Common Autoimmune Diseases. Immunity, 52 (6), 1119–1132.e1114. https://doi.org/10.1016/j.immuni.2020.04.006

Orrù, V., Steri, M., Sidore, C., Marongiu, M., Serra, V., Olla, S., Sole, G., Lai, S., Dei, M., Mulas, A., Virdis, F., Piras, M. G., Lobina, M., Pitzalis, M., Deidda, F., Loizedda, A., Onano, S., Zoledziewska, M., Sawcer, S., Devoto, M., Gorospe, M., Abecasis, G. R., Floris, M., Pala, M., Schlessinger, D., Fiorillo, E., & Cucca, F. (2020). Complex genetic signatures in immune cells underlie autoimmunity and inform therapy. Nat Genet, 52 (10), 1036–1045. https://doi.org/10.1038/s41588-020-0684-4

Park, A., Oh, S., Jung, K. L., Choi, U. Y., Lee, H. R., Rosenfeld, M. G., & Jung, J. U. (2020). Global epigenomic analysis of KSHV-infected primary effusion lymphoma identifies functional. Proc Natl Acad Sci U S A, 117 (35), 21618–21627. https://doi.org/10.1073/pnas.1922216117

Qian, J., Wang, Q., Dose, M., Pruett, N., Kieffer-Kwon, K. R., Resch, W., Liang, G., Tang, Z., Mathé, E., Benner, C., Dubois, W., Nelson, S., Vian, L., Oliveira, T. Y., Jankovic, M., Hakim, O., Gazumyan, A., Pavri, R., Awasthi, P., Song, B., Liu, G., Chen, L., Zhu, S., Feigenbaum, L., Staudt, L., Murre, C., Ruan, Y., Robbiani, D. F., Pan-Hammarström, Q., Nussenzweig, M. C., & Casellas, R. (2014). B cell super-enhancers and regulatory clusters recruit AID tumorigenic activity. Cell, 159 (7), 1524–1537. https://doi.org/10.1016/j.cell.2014.11.013

Ricigliano, V. A., Handel, A. E., Sandve, G. K., Annibali, V., Ristori, G., Mechelli, R., Cader, M. Z., & Salvetti, M. (2015). EBNA2 binds to genomic intervals associated with multiple sclerosis and overlaps with vitamin D receptor occupancy. PLoS One, 10 (4), e0119605. https://doi.org/10.1371/journal.pone.0119605

Ristori, G., Cannoni, S., Stazi, M. A., Vanacore, N., Cotichini, R., Alfò, M., Pugliatti, M., Sotgiu, S., Solaro, C., Bomprezzi, R., Di Giovanni, S., Figà Talamanca, L., Nisticò, L., Fagnani, C., Neale, M. C., Cascino, I., Giorgi, G., Battaglia, M. A., Buttinelli, C., Tosi, R., & Salvetti, M. (2006). Multiple sclerosis in twins from continental Italy and Sardinia: a nationwide study. Ann Neurol, 59 (1), 27–34. https://doi.org/10.1002/ana.20683

Sabari, B. R., Dall’Agnese, A., Boija, A., Klein, I. A., Coffey, E. L., Shrinivas, K., Abraham, B. J., Hannett, N. M., Zamudio, A. V., Manteiga, J. C., Li, C. H., Guo, Y. E., Day, D. S., Schuijers, J., Vasile, E., Malik, S., Hnisz, D., Lee, T. I., Cisse, I. I., Roeder, R. G., Sharp, P. A., Chakraborty, A. K., & Young, R. A. (2018). Coactivator condensation at super-enhancers links phase separation and gene control. Science, 361 (6400). https://doi.org/10.1126/science.aar3958

Schwalb, B., Michel, M., Zacher, B., Frühauf, K., Demel, C., Tresch, A., Gagneur, J., & Cramer, P. (2016). TT-seq maps the human transient transcriptome. Science, 352 (6290), 1225–1228. https://doi.org/10.1126/science.aad9841

Sheffield, N. C., & Bock, C. (2016). LOLA: enrichment analysis for genomic region sets and regulatory elements in R and Bioconductor. Bioinformatics, 32 (4), 587–589. https://doi.org/10.1093/bioinformatics/btv612

Simeonov, D. R., Gowen, B. G., Boontanrart, M., Roth, T. L., Gagnon, J. D., Mumbach, M. R., Satpathy, A. T., Lee, Y., Bray, N. L., Chan, A. Y., Lituiev, D. S., Nguyen, M. L., Gate, R. E., Subramaniam, M., Li, Z., Woo, J. M., Mitros, T., Ray, G. J., Curie, G. L., Naddaf, N., Chu, J. S., Ma, H., Boyer, E., Van Gool, F., Huang, H., Liu, R., Tobin, V. R., Schumann, K., Daly, M. J., Farh, K. K., Ansel, K. M., Ye, C. J., Greenleaf, W. J., Anderson, M. S., Bluestone, J. A., Chang, H. Y., Corn, J. E., & Marson, A. (2017). Discovery of stimulation-responsive immune enhancers with CRISPR activation. Nature, 549 (7670), 111–115. https://doi.org/10.1038/nature23875

Sloan, C. A., Chan, E. T., Davidson, J. M., Malladi, V. S., Strattan, J. S., Hitz, B. C., Gabdank, I., Narayanan, A. K., Ho, M., Lee, B. T., Rowe, L. D., Dreszer, T. R., Roe, G., Podduturi, N. R., Tanaka, F., Hong, E. L., & Cherry, J. M. (2016). ENCODE data at the ENCODE portal. Nucleic Acids Res, 44 (D1), D726–732. https://doi.org/10.1093/nar/gkv1160

Sun, Y., Peng, I., Senger, K., Hamidzadeh, K., Reichelt, M., Baca, M., Yeh, R., Lorenzo, M. N., Sebrell, A., Dela Cruz, C., Tam, L., Corpuz, R., Wu, J., Sai, T., Roose-Girma, M., Warming, S., Balazs, M., Gonzalez, L. C., Caplazi, P., Martin, F., Devoss, J., & Zarrin, A. A. (2013). Critical role of activation induced cytidine deaminase in experimental autoimmune encephalomyelitis. Autoimmunity, 46 (2), 157–167. https://doi.org/10.3109/08916934.2012.750301

Umeton R., S. G., Sorathiya A., Liò P., Papini A., and Nicosia G. (2011). Design of robust metabolic pathways. In: In Proceedings of the 48th Design Automation Conference (DAC ’11). ACM, New York, NY, USA, 747–752.

Vahedi, G., Kanno, Y., Furumoto, Y., Jiang, K., Parker, S. C., Erdos, M. R., Davis, S. R., Roychoudhuri, R., Restifo, N. P., Gadina, M., Tang, Z., Ruan, Y., Collins, F. S., Sartorelli, V., & O’Shea, J. J. (2015). Super-enhancers delineate disease-associated regulatory nodes in T cells. Nature, 520 (7548), 558–562. https://doi.org/10.1038/nature14154

van Arensbergen, J., Pagie, L., FitzPatrick, V. D., de Haas, M., Baltissen, M. P., Comoglio, F., van der Weide, R. H., Teunissen, H., Võsa, U., Franke, L., de Wit, E., Vermeulen, M., Bussemaker, H. J., & van Steensel, B. (2019). High-throughput identification of human SNPs affecting regulatory element activity. Nat Genet, 51 (7), 1160–1169. https://doi.org/10.1038/s41588-019-0455-2

Villamil, G., Wachutka, L., Cramer, P., Gagneur, J., & Schwalb, B. (2019). Transient transcriptome sequencing: computational pipeline to quantify genome-wide RNA kinetic parameters and transcriptional enhancer activity. bioRxiv, 659912. https://doi.org/10.1101/659912

Weinert, B. T., Narita, T., Satpathy, S., Srinivasan, B., Hansen, B. K., Schölz, C., Hamilton, W. B., Zucconi, B. E., Wang, W. W., Liu, W. R., Brickman, J. M., Kesicki, E. A., Lai, A., Bromberg, K. D., Cole, P. A., & Choudhary, C. (2018). Time-Resolved Analysis Reveals Rapid Dynamics and Broad Scope of the CBP/p300 Acetylome. Cell, 174 (1), 231–244.e212. https://doi.org/10.1016/j.cell.2018.04.033

Wilson, D. J. (2019). The harmonic mean. Proc Natl Acad Sci U S A, 116 (4), 1195–1200. https://doi.org/10.1073/pnas.1814092116

Wong, E. S., Zheng, D., Tan, S. Z., Bower, N. L., Garside, V., Vanwalleghem, G., Gaiti, F., Scott, E., Hogan, B. M., Kikuchi, K., McGlinn, E., Francois, M., & Degnan, B. M. (2020). Deep conservation of the enhancer regulatory code in animals. Science, 370 (6517). https://doi.org/10.1126/science.aax8137

