## Supplementary Figure 1 for "GWAS associated Variants, Non-genetic Factors, and Transient Transcriptome in Multiple Sclerosis Etiopathogenesis: a Colocalization Analysis"

**Figure Supplement**

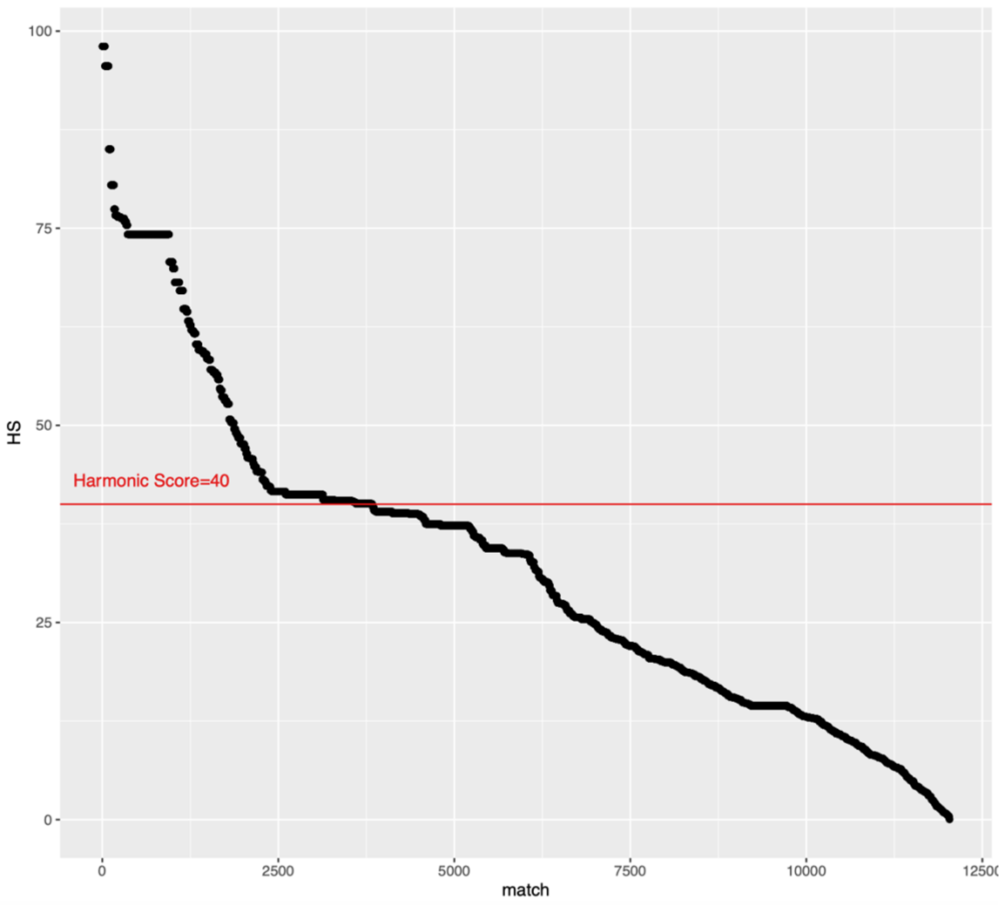


**Figure Supplement 1. Harmonic Score threshold defining the top colocalization hits.** The plot shows all of colocalization matches, ranked by Harmonic Score (HS). The curve inflection point, highlighted with a red line, suggests a threshold for selecting the most relevant hits (i.e., those scoring at HS>40).
